# Dopamine enhances population coding of fear in prefrontal cortex

**DOI:** 10.1101/2023.01.17.524376

**Authors:** Jeremy Lesas, Feugas Pierre, Cyril Herry, Cyril Dejean

## Abstract

Fear acquisition is a survival strategy enabling the prediction of potential threat on the basis of reliable environmental cues identified through associative learning. In this study we investigated the role of dorsomedial prefrontal cortex (dmPFC) – ventral tegmental area (VTA) coordination in fear acquisition. Despite the implication of a VTA-dmPFC circuit in fear acquisition, the mechanism by which the two structures interact and, most particularly, the manner in which dmPFC neuronal dynamics are impacted by DA remain largely unexplored. The present paper addresses this question by means of behavioral, optogenetic, and electrophysiological experiments. We first characterize the dynamics of VTA neurons and their relation with dmPFC during fear acquisition to then manipulate this dopaminergic inputs to dmPFC and address its role in fear memory expression and its dmPFC neuronal correlates. Our data first unravel that VTA neurons develop a conditioned stimulus coding activity through consecutive pairings with the unconditioned aversive stimulus and maintain this predictive activity overnight. Shutting down these VTA inputs to dmPFC resulted in a reduction in fear behavior and changes in population coding. These alteration were specific to spontaneous expression of fear and the population coding supporting that behavior. Our data show that dopamine role is to increase the accuracy of fear coding in dmPFC.

## Introduction

Fear acquisition is a survival strategy enabling the prediction of potential threat on the basis of reliable environmental cues identified through associative learning ^1^. The dorsomedial prefrontal cortex (dmPFC) has been identified as a key brain structure for the acquisition and the expression of associative fear behaviour^2–4^ and interacts in a synchronous manner with other brain structures involved in fear behaviour to elicit a variety of measurable behavioural outputs ^5^. Among these structures, the ventral tegmental area (VTA) is a central actor in associative learning as a part of the dopaminergic system. It mediates learning through predictive temporal association of outcome-value and value-predicting stimuli ^6,7^. Although the bulk of VTA DA neurons are preferentially activated by appetitive outcome stimuli, seminal papers also identified a subset of neurons excited by aversive stimuli^6,8,10–12^ and recent research points to a necessary role for DA in fear learning ^9^.

Despite the implication of a VTA-dmPFC circuit in fear acquisition, the mechanism by which the two structures interact and, most particularly, the manner in which dmPFC neuronal dynamics are impacted by DA remain largely unexplored, although related research in the fear domain provides solid groundwork. Indeed, two mechanisms are currently thought to underpin associative fear behavior. The first one consists in the disihnibition and theta-band (8-12Hz) oscillatory synchronization of dmPFC neurons during conditioned stimulus (CS) presentation^13^. Both during and between CS, a second mechanisms consists in dmPFC neurons synchronization into assemblies that temporally control fear behaviour^14^. During fear expression, these assemblies phase lock to oscillations centered on 4 Hz (3 – 6 Hz), to such an extent that induction of 4Hz oscillations in dmPFC neurons can elicit both the learning and expression of fear behaviour^15^ prefrontal-amygdala circuits also synchronize at 4 Hz during fear expression^15^. Interestingly, a 4 Hz coordination of prefrontal cortex-ventral tegmental area circuits has been observed during reward based decision making^16^ indicating the possibility that a similar oscillatory synchronization of the VTA and dmPFC may play a role in fear acquisition. Also, the emergence of neuronal assemblies through increased pairwise oscillatory synchronization within prefrontal cortex neurons is influenced by DA^17^ the probable source of which is the VTA. Overall, the literature supports neuronal oscillatory synchronization as a key mechanism to both fear learning and expression and suggests that cortical synchronization in general is influenced by DA release from mesocortical DA projections.

Yet the role of mesocortical DA in fear related neuronal synchronization in dmPFC remains unclear. The present paper addresses this question by means of behavioral, optogenetic, and electrophysiological experiments. We first characterize the dynamics of VTA neurons and their relation with dmPFC during fear acquisition to then manipulate this dopaminergic inputs to dmPFC and address its role in fear memory expression and its dmPFC neuronal correlates.

## Methods

### Animals

Male C57BL6/J or DAT-Cre C57BL6/J mice (8-11 weeks old) were individually housed for at least 7 days before all experiments, under a 12 hour light–dark cycle, and provided with food and water *ad libitum*. All procedures were performed in accordance with standard ethical guidelines (European Communities Directive 86/60-EEC) and were approved by the committee on Animal Health and Care of Institut National de la Santé et de la Recherche Médicale and French Ministry of Agriculture and Forestry (authorization A3312001).

### Material and surgeries

#### Wild Type animals

Wild type animals underwent a single surgery for electrode implantation. Mice were anaesthetized with isoflurane (induction 3%, maintenance 1.5%) in oxygen. Body temperature was maintained at 37° C with a temperature controller system. Mice were secured in a stereotaxic frame where analgesia was provided by local subcutaneous injection of lidocaine and intraperitoneal injection of meloxicam. Mice were unilaterally implanted with 2 electrode arrays in the left dmPFC and left VTA (dmPFC coordinates: 2 mm anterior to bregma; 0. 4 mm lateral to midline; 1.4 ventral to cortical surface; VTA coordinates: at the following coordinates: 3.52 mm posterior to bregma; 1.3 mm lateral to midline; 4 mm ventral to cortical surface; angle of approach 15°.; The wires were attached to two 18-pin connectors (Omnetics). All implants were secured using Super-Bond cement (SunMedical). After surgery, mice were allowed to recover for at least 7 days and were habituated to handling before entering the behavioural experiment.

#### DAT-Cre animals

Transgenic mice underwent two surgeries: an initial viral injection surgery and subsequent optic fiber and optrode implantation. Mice were anaesthetized with isoflurane (induction 3%, maintenance 1.5%) in oxygen. Body temperature was maintained at 37° C with a temperature controller system. Mice were secured in a stereotaxic frame where analgesia was provided by local subcutaneous injection of lidocaine and intraperitoneal injection of meloxicam. To enable optical control of midbrain DA neurons, test mice (*n* = 3) were bilaterally injected with conditional AAV encoding Cre-dependent ArchT (rAAV9/CAG-Flex-ArchT-GFP, 0.2 – 0.25 μl per hemisphere) using glass micropipettes (tip diameter: 10 – 20 μm) at the following coordinates: 3.52 mm posterior to bregma; 1.3 mm lateral to midline; 4 mm ventral to cortical surface; angle of approach 15°. Control mice (*n* = 4) were injected with similar bilateral quantities of AAVs expressing only GFP. 10 – 12 days after the injection, mice were unilaterally implanted with an optic fiber in the left dorsomedial prefrontal cortex (dmPFC) and an optrode in the right dmPFC at the following coordinates: 2 mm anterior to bregma; 0.65 mm lateral to midline; 1.45 ventral to cortical surface; angle of approach 15°. Optic fibers (Doric Lenses) had a diameter of 200 μm, numerical aperture of 0.37, and a flat tip. Optrodes consisted of 32 individually insulated, gold-plated nichrome wires (13 μm inner diameter, impedance 30–100 KV; Kanthal) positioned around an optic fiber. The wires were attached to two 18-pin connectors (Omnetics). Optic fibers and optrodes were assembled manually. All implants were secured using Super-Bond cement (SunMedical). After surgery, mice were allowed to recover for at least 7 days and were habituated to handling. For optimal virus expression prior to optogenetic intervention, all experiments were performed 3-5 weeks post-injection.

### Behavioral paradigm

Fear conditioning and testing were conducted in distinct contexts (context A and context B). Context A was cleaned with 70% ethanol and context B with 1% acetic acid before and after each session. To score freezing behaviour, an automated infrared beam detection system located on the bottom of the experimental chamber was used (Imetronic). The animals were considered to be freezing if no movement was detected for 2 seconds. On day 1, DAT-Cre C57BL6/J mice were submitted to an habituation session in context A in which they received four presentations of the CS^+^ and of the CS^−^ (total CS duration, 30□s; consisting of 50-ms pips at 0.9 Hz repeated 27 times, 2 ms rise and fall; pip frequency, 7.5 kHz or white-noise, 80 dB sound pressure level). Discriminative fear conditioning was performed in a subsequent session 2 hours later on the same day by pairing the CS^+^ with a US (1 s foot-shock, 0.6 mA, 5 CS^+^–US pairings; inter-trial intervals, 20–180 s). The onset of the US coincided with the offset of the CS^+^. The CS^−^ was presented after each CS^+^/US association but was never reinforced (five CS^−^ presentations; inter-trial intervals 30-180 s). On day 2, conditioned mice were submitted to a test session in context B during which they received 4 presentations of the CS^−^ followed by 4 presentations of the CS^+^ (total CS duration 30 s; consisting of 50 ms pips at 0.9 Hz repeated 27 times, 2 ms rise and fall; pip frequency 7.5 kHz (CS^+^) or white-noise (CS^−^), 80 dB sound pressure level. During all experimental sessions, optrode connectors were connected to two headstages (Plexon) comprised of sixteen unity-gain operational amplifiers. Headstages were connected to a 32-channel preamplifier (gain 100X bandpass filter from 150 Hz to 9 kHz for unit activity and from 0.7 Hz to 170 Hz for field potentials, Plexon). Both basic optic fibers and optrodes were connected to a laser (OptoDuet 593 nm, Ikecool) via fiber-optic patch cords (diameter 200 μm, Doric Lenses). Both the headstage and optic fiber connections were fixed to rotary equipment which allowed the mice to move freely in the behavioral chamber.

### Optogenetics and recordings

During the conditioning session, mice received bilateral yellow light stimulations (approx. 17 mW, 593.5 nm, 5 ms pulses repeated at 20 Hz for 30 s) during presentation of CS^+^ and subsequent US in order to inhibit the terminals of ArchT-expressing VTA-dmPFC projecting DA neurons. The rationale behind this was derived from studies on the dynamic nature of dopaminergic activity during associative learning (Schultz 1998 for review) showing that a spontaneous DA release upon unexpected occurrence of emotionally valent events (reward or, here, aversive event, i.e. US) temporally shifts during learning leading to a DA release peak upon presentation of event-predicting stimulus (e.g. CS^+^), in keeping with classical Pavlovian conditioning models. Control GFP-expressing mice were submitted to the same protocol to control for any behavioral modification that may occur as a direct result of internal light stimulation effects (illumination, heating, etc.). If inhibition of DA release during conditioning has an effect on associative fear learning, it should be measurable through comparison of freezing in the ArchT population and GFP population during the postconditioning test session. Single unit activity was recorded during habituation, conditioning, and test sessions for the optrode implanted ArchT mice (*n* = 3) and for habituation and test sessions for the GFP (control) mice (*n* = 4). An additional 2 optrode implanted, ArchT injected mice were excluded from analysis as they displayed extremely high generalized anxiety and immobility even in habituation sessions, meaning it was not possible to distinguish conditioned fear expression from general stress-induced fear-like behaviour.

### Histology and immunohistochemistry

Mice were euthanized with isoflurane and perfused through the left ventricle, first with 1% NaCl and then with a 4% paraformaldehyde solution (PFA). The brains were extracted and post-fixed in PFA for 24 hours. For immunostaining and histological verification of implantation sites, 40 μm coronal brain sections were made using a vibratome and were stored in sodium-azide PBS. To assess the specificity of AAV infection of dopaminergic neurons, immunohistochemistry with antibodies against tyrosine hydroxylase was performed. Brain sections were rinsed in PBS three times and then incubated in a blocking solution containing 5% donkey serum and 0.3% triton for 1.5 hours. Brain sections were then separated into PFC sections and VTA sections. PFC sections were incubated overnight at 4°C in a blocking solution containing primary antibodies against green fluorescent protein (GFP; 1:1000 dilution, raised in goat, AB6673, Abcam). VTA sections were incubated overnight in a blocking solution containing antibodies for green fluorescent protein (GFP; 1:1000 dilution, raised in goat, AB6673, Abcam) and tyrosine hydroxylase (TH; 1:1000 dilution, raised in mouse, MAB318, Millipore). Brain sections were again rinsed in PBS three times. Sections were then incubated in secondary antibodies for at least 1.5 hours: PFC sections in Alexa Fluor 488-conjugated donkey anti-goat antibody (1:1000 dilution, A11055, Molecular Probes); VTA sections in Alexa Fluor 488-conjugated donkey anti-goat antibody (1:1000 dilution, A11055, Molecular Probes) and Alexa Fluor 594-conjugated donkey antimouse antibody (1:100 dilution, A21203, Molecular Probes). Sections were given a final three rinses in PBS before being mounted onto glass slides with VECTASHIELD, coverslip, and stored in the dark at 4°C. Optic fiber and optrode location accuracy was assessed visually by the cavity left in the cortex by the fibers.

### Data acquisition and analysis

Freezing behavior was measured using the aforementioned automated infrared beam detection system (Imetronic) and collected through the PolyFear software. For each session, this data was analyzed and temporally correlated to presentations of CS^−^ and CS^+^ using custom scripts written in Matlab. Spiking activity collected through the headstages was digitized at 40 kHz, bandpass filtered from 250 Hz to 8 kHz, and isolated by time-amplitude window discrimination and template matching using a Multichannel Acquisition Processor system (Plexon). Single-unit spike sorting was performed using Off-Line Spike Sorter (OFSS, Plexon) for all behavioral sessions. Principal component (PC) scores were calculated for each waveform and were plotted in a 3D PC space. This presentation facilitated the manual selection of waveform clusters. Multiple waveforms were classified as those coming from a single neuron if they formed a discrete and isolated cluster in the PC space and if their autocorrelograms displayed a clear refractory period of at least 1 ms. To avoid analysis of the same neuron recorded on different channels, cross-correlation histograms were calculated with a window width of 1 s (−500 ms to 500ms) and a bin size of 5 ms. If a target neuron presented a peak of activity at the same time that the reference neuron fired, only one of the two neurons was considered for further analysis. These validated cross-correlograms (CC) were analyzed in order to identify pairwise synchronization between all recorded neurons for each animal. Moment to moment variations in each CC were quantified using a Z-score normalization, thresholding of the basal cross-correlogram mean, and standard deviation calculation using the −500 to −250 ms portion of the CC. Only those neurons whose CC peak lay outside 1.65 standard deviations (alpha = 0.05, unilateral test) were retained as significantly modulated. Beyond moment to moment variation, it is possible that the inner variability between a pair of firing neurons may produce significant deflection in the CC. To account for this in the case of every pair of reference and target neurons, the reference spike train was shuffled and its corresponding CC calculated. This shuffling procedure was repeated 25 times for each reference neuron. We then used the mean of this surrogate data set to establish a confidence limit of 95% for assessing the significance of CC variability (alpha = 0.05, unilateral test). A CC crossing the upper limit of both the Z-score and the surrogate confidence interval indicated that reference and target neurons presented significant co-activation.

We investigated predictive power of network activity for fear behaviour. To this end we analysed network state as the joint activity of all simultaneously recorded dmPFC principal cells. For n PNs simultaneously recorded in a session of L milliseconds length, single unit rate histograms were first calculated with a sliding window of 150 ms and (50 ms overlap) then concatenated in the neuron dimension returning a n by L/50 matrix. In this study we were interested in labelling which neurons were co-activated rather than how much they were co-activated. We therefore focused on the sole presence or absence of activity for a neuron inside each time bin independently of its actual firing rate. To this end, the rate histogram matrix was binarized such that any bin value strictly above zero was given a value of 1 (otherwise 0). Note that other type of normalization of firing rates such as z-scores yielded very comparable results. In order to retrieve not only the activity but also the joint activity of the n neuron population we calculated the n by n correlation matrix (CM) of the population as follow. Considering the ith and jth PN in a single time bin t, if PN(i,t) and PN(j,t) are both active CM(i,j,t) is given a correlation score of 1. In any other case, CM(i,j,t) is given a score of 0. This calculation is applied for all neuron pairs inside each time bin, hence returning a CM of n by n by L/50 the captured the instantaneous correlation profiles for the whole PN population along the time axis. To investigate the emergence of specific network state during the recording session a principal component analysis was performed on the CM. The score on the 1st principal component (PC1) was our proxy for network state. The probability of correct freezing prediction by PC score was then retrieved by quantifying the amount of true and false positive in a ROC analysis. Same method was used with sets of surrogate spike trains to define hazard prediction power and its variability.

### Local field potential analyses

Power spectral density (PSD) for CS^+^ periods was retrieved by averaging the results of the time-resolved Fourier analysis across this specific time interval. To average LFP power across animals, we first normalized the PSD histogram as the percentage of total power between 3 and 16 Hz. We then compared ArchT and GFP average power in the 3 to 6 Hz band, encompassing the central 4 Hz frequency.

In order to analyze the spike-phase relationship, local field potentials recorded from each animal and for each session were filtered using a 3-6 Hz (i.e. the 4 Hz oscillations range) bandpass filter. As it had already been shown that neurons in the mPFC involved in fear expression are preferentially phase locked to the ascending phase of 4 Hz oscillations, we calculated the instantaneous phase of the 3-6 Hz filtered LFP using the Hilbert transform in order to see which of our recorded neurons fell under this criteria. Each spike of each neuron was assigned its corresponding 4 Hz phase value and, from this, we were able to construct phase histograms for each neuron using bins of 36°. Phase locking significance was then assessed from these histograms using Rayleigh’s test for circular uniformity (alpha = 0.05, bilateral test).

## RESULTS

### VTA neurons develop fear coding during Pavlovian conditioning

In order to evaluate the firing behavior of VTA neurons along fear conditioning, mice were chronically implanted with recording electrodes targeting the VTA, and submitted to a discriminative fear conditioning procedure (Fig. 1a). In this learning paradigm, mice learn to discriminate between two auditory stimuli of different frequencies. The first CS (CS+) is associated with the delivery of a coincident mild foot-shock (the US) whereas the second one (the CS−) is not. Twenty-four hours following fear conditioning, when reexposed to the CS+ but not the CS−, mice displayed a selective increase in conditioned freezing behaviour, a characteristic fear immobilization reaction (Fig. 1a). After this verification that all 7 animals had properly associated CS+ to the footshock, examination of their learning curve during conditioning revealed a gradual increase in freezing behavior from the first CS+ presentation to the fifth and last one (Fig. 1a). In order to assess a putative change in VTA neuron CS+ coding along CS− US pairings we discriminated early (CS #1 & #2) and late pairings (CS #4 & #5) and compared the response of VTA putative principal neurons (PN, extended data Fig. 1) to CS+ presentations (pips) for both epochs (Fig. 1b-d). Among 81 cells, we found that several VTA PNs acquired enhanced response to late vs. early CS+. We identified 2 types of enhancement with one subset of neurons showing an absolute increase in post-pip firing (dynamic rate code, 13%, Fig. 1b, extended data Fig. 2a) and another subset showing an increase relative to pre-pip activity for late CS that was not detected with early CS+ (dynamic signal to noise code, 10%, Fig. 1c, extended data Fig. 2a). Hence along CS− US pairings, 23% of VTA PN develop a significant response to consecutive CS that mirrors the gradual acquisition of associative fear behavior during learning (Fig. 1a). Dynamic rate and signal to noise coders were more likely to respond to the shock than other cells and did it in a significantly higher magnitude (Fig. 1d, extended data Fig. 2b). On test day (Fig. 2), those neurons that developed CS coding retained this functional feature as cell that were considered to be conserved under our electrodes overnight displayed a strong phasic activation at CS presentation (Fig. 2e, *top*). Very interestingly, these cells showed a marked suppression in firing at the time of US omission by the end of the CS (Fig. 2e, *top*). Because VTA PN are projection neurons, this acquired fear related activity is likely to reach their distant targets and participate to the acquisition of neuronal coding in the former.

**Figure 01:**
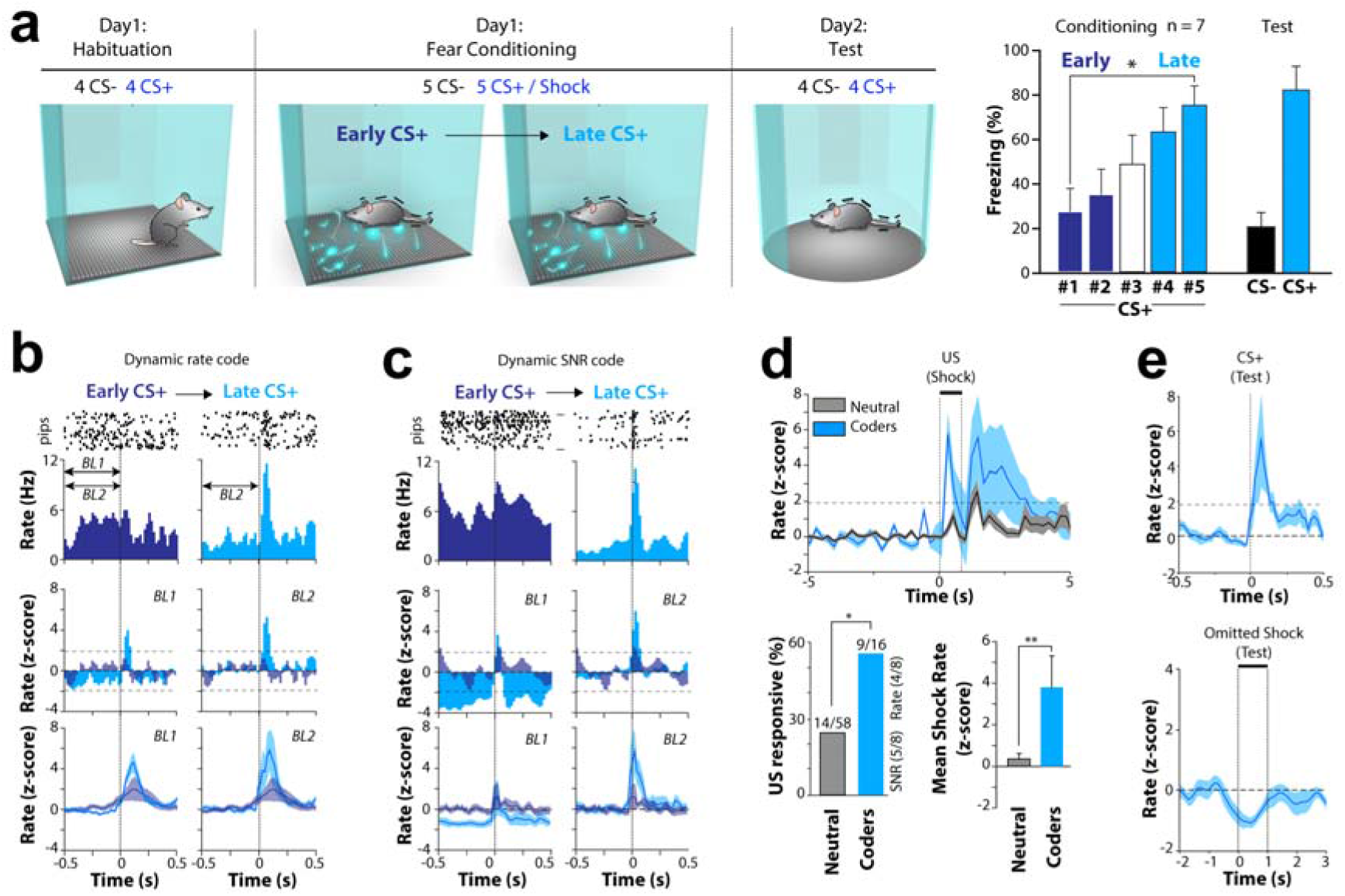
VTA neurons develop fear coding during Pavlovian conditioning. a. Experimental paradigm (left) and behavioural results (right). Based on the former, the analysis of neuronal coding for CS during conditioning was splitted into early and late CS+ presentations (n = 7, one-way repeated measures ANOVA, F(6,4) = 4.372, p = 0.009, Bonferroni post-hoc test: CS#1 vs CS#5, p = 0.015). b-d. Development of two neuronal codes for CS+ between early and late CS presentations. b. Top. Example of a neuron acquiring a dynamic rate code for CS+ as displayed by raster plot and corresponding PSTH. Middle. Neuronal response to the CS normalized to BL1 (left) and BL2 (right) shows this cell is not responsive to early CS+ as shown on the raster plot (top) and the PSTH (bottom). Following late CS+ pips the cell develops a dynamic increase in firing rate (z-score > 1.96 for 3 consecutive 20ms bins p < 0.05). Bottom. Averaged PSTH for neurons acquiring dynamic rate code. c. Example of a neuron acquiring a dynamic signal-to-noise (SNR) code for CS+. Top. This cell is not responsive to early CS+ as shown on the raster plot and the PSTH. During late CS+, the cell does not increase firing rate in response to the sound compared to early baseline (BL1 normalization, Middle Left) but display a decrease in tonic firing around pip occurrence. Normalizing by late pre-CS+ activity (BL2, Middle Right) reveals a dynamic increase in firing rate in response to the sound (blue trace, zscore >1.96 for 3 consecutive bins, p < 0.05). These cells code for CS+ through an increase in Signal to noise ratio rather than a raw increase in firing rate. Bottom. Averaged PSTH for neurons acquiring dynamic signal-to-noise. d. Most neurons acquiring coding for CS+ also respond to the shock. Top. Average Peri-Shock time histogram for the 18 neurons acquiring coding for CS+ along condition. Bottom. Proportion (Left) and magnitude (Right) of neuronal activation during the shock among neutral and CS+ coder cells (Proportions: Chi^2^(1)=4.631; p=0.031; Magnitude: Mann-Whitney test: T = 851,000 n(small)= 16 n(big)= 60, p = 0,003). e. On day 2 during Test, coding neurons respond to CS+ and shock omission. Top: Average peri-CS+ time histogram for the 10 neurons acquiring coding for CS+ along condition that could be recorded on Day2 during test session. Bottom: Average peri-shock time histogram for the same 10 neurons. These neurons show a tendency to be suppressed at the time the shock is supposed to come while the majority of them where activated at the time of the shock. Shaded areas: mean +/− SEM.

**Figure 02:**
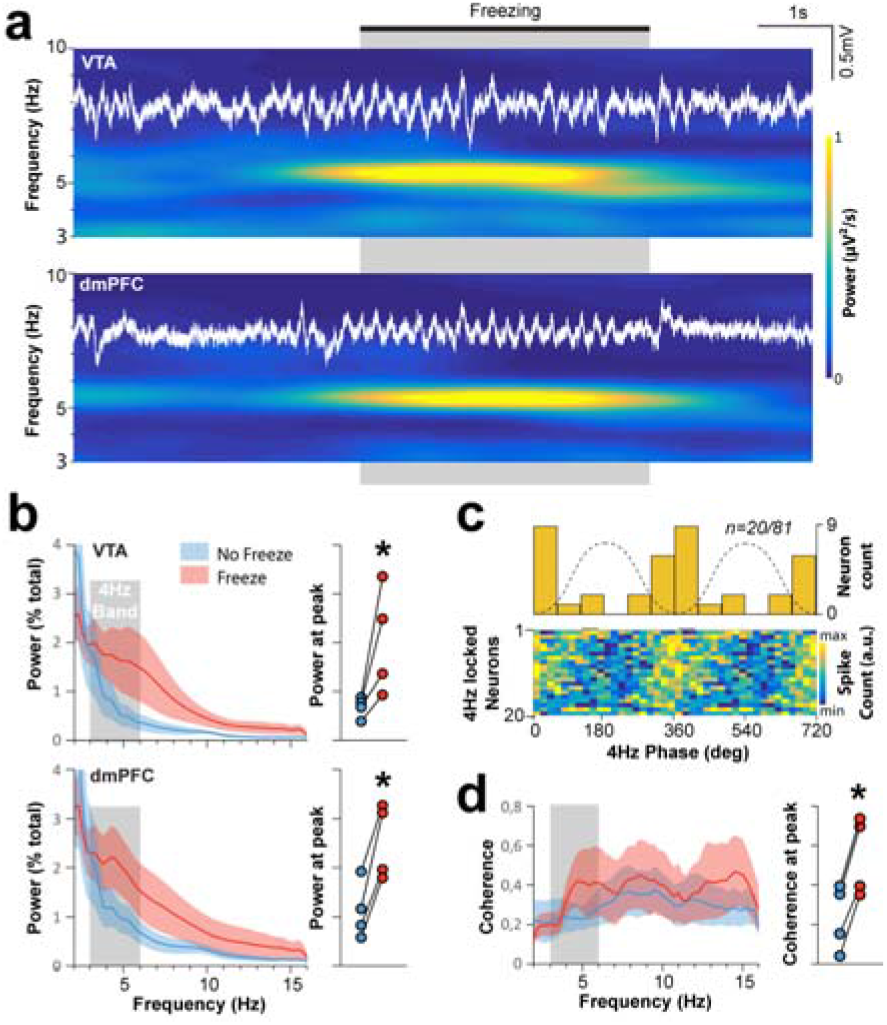
VTA and dmPFC oscillatory synchronization during fear conditioning. During conditioning, VTA and dmPFC LFP 4Hz power increases when animals express fear. a. Example raw VTA LFP around a freezing epoch and the corresponding spectrogram displaying power as a function of time and frequency (color scale on the right). b. Left. Power spectral density plot, displaying the change in VTA LFP power in the 4Hz band between freezing and non-freezing epochs (n = 11). Right. Distribution of mean power ratio between 3-7Hz (Wilcoxon test. VTA: Z(10) = −2.934, *** p < 0.001) c. VTA neurons significantly phase lock to 4Hz LFP oscillations when the animals freeze during conditioning (n = 20/81, 24%)). Note that neurons acquiring coding for CS+ present a higher phase-locking probability (61%, n=11/18) than neutral neurons (14.5%, n=9/63, Chi^2^(1)=13.763, p<0.001). d-f. Example for dmPDC LFP and neurons. e. dmPFC: Z(6) = −2.366, * p = 0.016. f. dmPFC neurons (n = 16/65, 25%) g. Cartoon of the scheme of simultaneous VTA-dmPFC recordings in animals undergoing Pavlovian auditory fear conditioning. h. During conditioning, VTA and dmPFC preferentially synchronize around 4Hz during freezing. Left. Coherence plot, measuring the level of synchronization between VTA and dmPFC LFP oscillation as a function on frequency (n =7). Right. Distribution of coherence ratio (Freezing/NoFreezing) at peak coherence (Wilcoxon test: Z(6) = −2.366, * p = 0.016) Shaded areas: mean +/− SEM.

### VTA and dmPFC oscillatory synchronization during fear conditioning

Given the well documented implication of dmPFC in fear behavior (^5^) and its position as a preferred target for VTA neurons involved in fear acquisition (^11^), we investigated their possible joint activity during fear conditioning in the same paradigm as precedently. Each structure presented an increase in 4Hz power during freezing (Fig. 2a,b d & e). Spike-phase analysis showed that a substantial amount of PNs significantly lock onto that rhythm (VTA: 24%, dmPFC: 25%), showing that the presence of 4 Hz oscillations in the LFP of both structures reflects neuronal activity and discards the possibility that it is solely due to passive conduction. Importantly, the proportion of phase-locked neurons was significantly higher in CS+ coding neurons than in the rest of VTA population (Fig. 2c). Finally we analyzed VTA-dmPFC synchronization in mice chronically implanted with recording electrodes targeting both the VTA and the dmPFC (Fig. 2g). In these animals, coherence analysis during fear conditioning revealed that VTA and dmPFC LFPs synchronize preferentially between 3 and 6Hz (4Hz band) during freezing episodes (Fig. 2h). These results confirm the entrainment of dmPFC by 4Hz rhythm during fear conditioning, unravels the same process for VTA and suggests a functional interaction between VTA and dmPFC during associative fear conditioning.

### Dopaminergic projection from VTA to dmPFC modulates fear learning

To address the causal role of VTA to dmPFC projection in fear learning we combined behavioural, electrophysiological and optogenetics techniques. Based on last paragraph results, we designed a two-step experimental strategy (Fig. 3a). On day 1 during fear conditioning, we suppressed dopaminergic inputs to dmPFC at the time of CS+ successive presentations (step 1). On day 2 during test, we assessed VTA inputs suppression on fear behaviour and dmPFC coding mechanisms (step 2). To this end, we injected the VTA of 12 animals with a floxed adeno associated virus carrying ArchT and GFP (n=7) or GFP only (n=5). In order to inhibit specifically VTA inputs to dmPFC (step1) and then record dmPFC electrophysiological activity *in vivo* (step2), we implanted an optrode in the dmPFC to illuminate DA terminals and record dmPFC activity (Fig. 3b). Visual microscope inspection after immunohistochemistry confirmed that all ArchT animals had been successfully infected with the AAV with a consistent overlap of GFP with dopaminergic cell bodies (Supplementary Fig. 3) and GFP labeled projections in the dmPFC (Fig. 3d) where optrodes had been implanted (Supplementary Fig. 3). During conditioning, light stimulations induced a delay in the learning curve of ArchT group animals compared to that of the GFP control group, although both groups reached similar freezing levels by the end of conditioning (Fig. 3c). This acute effect of dopamine suppression in dmPFC validates our methodology and brings a crucial requisite in our strategy. Moving to step 2, on test day (Fig. 3d-e) both group displayed similar freezing levels to CS+ (high) compared to CS− (low) indicating that discriminative fear learning was achieved despite optogenetic manipulation. Nonetheless, we observed a consistent decrease in fear expression in-between CS+ presentations intervals (*outside CS+*, Fig. 3e) where most freezing behaviour usually occurs in this paradigm^14,15,18^ explained by a decrease in the amount of freezing episode count and therefore initiation (supplementary Fig. 4). Hence VTA dopaminergic inputs to dmPFC during fear conditioning support internally triggered and sustained fear expression but not CS− evoked recall of fear memory. Recent studies have evidenced two precise mechanisms underpinning CS− evoked fear^13,19^ on the one hand and internally sustained (or triggered) fear on the other^14,15^. To address the changes in dmPFC neuronal fear coding following the prior suppression of VTA inputs during FC, we analyzed LFP and single unit activity in ArchT and GFP animals.

**Figure 03:**
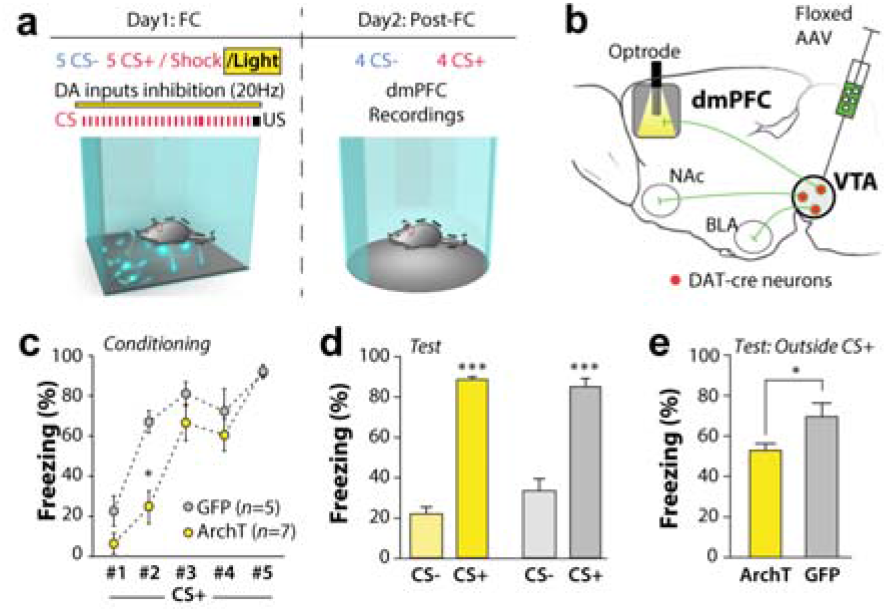
Dopaminergic projection from VTA to dmPFC modulates fear learning. a. Experimental scheme. b. Optogenetic strategy. c. learning curve. Animals which DA terminals were inhibited during CS− US pairings (ArchT group, yellow) display a delay in associative learning before eventually reaching the performance of GFP control animals (ArchT n = 7, GFP n = 5, Two-way ANOVA with one factor repetition, Factor 1: ArchT vs GFP, F(1,5) = 7.128, p = 0.024; Factor 2: epoch, F(1,2) = 42.785, p < 0.001; F1×F2, F(1,4) = 3.152, p = 0.024; Holm-Sidak post-hoc adjusted multiple comparison: within post-onset:, **p = 0.01; within CS2: *p < 0.001). d-e. Dopamine input silencing during conditioning decreases fear expression outside CS+ presentation during test. d. During memory testing on day 2, both ArchT and GFP animal discriminate between CS− and CS+. Freezing to CS+ was similar in both groups (ArchT n = 7, GFP n = 5, Two-way ANOVA with one factor repetition, Factor 1: ArchT vs GFP, F(1,8) = 3.093, p = 0.109; Factor 2: epochs, F(2,8) = 158.446, p < 0.001; F1×F2, F(1,1) = 2.268, p = 0.129; Holm-Sidak post-hoc adjusted multiple comparison: within ArchT, CS− vs CS+: ***p < 0.001; within GFP, CS− or CS+: ***p < 0.001). e. The magnitude of freezing outside CS+ was significantly decreased in ArchT animals (outside CS+: Mann-Whitney test, U = 5.00, *p = 0.048). Error bars: SEM.

### dmPFC coding mechanisms of conditioned stimulus is independent from previous dopaminergic modulation

CS− evoked fear relies on the disinhibition of dmPFC PNs that project to the basolateral amygdala, a mechanism which is also concurrent with post-pip resetting of local theta (8-12Hz) LFP oscillations^14^. To assess whether our manipulation impacted these mechanisms, we first investigated dmPFC PN responses to CS+ pips during test by means of peristimulus time histograms around CS+ occurrence. Neuronal responses to CS+ were similar in ArchT and GFP groups, whether it was in terms of shape (figure 4a,b) or in the proportions of neurons displaying positive, negative or no response (figure 4d). We next quantified post-pip LFP theta resetting in both group. This phenomenon implies that each pip is followed by an instantaneous stabilization of theta wave phase and the subsequent alignment of theta cycles from one pip to the next^21^. The mean resultant length (MRL) computed across pips gives an estimate of phase stability and we used MRL as an objective measure of oscillation resetting. ArchT and GFP groups showed post-pip LFP resetting very similar to each other and to that observed in aforementioned studies, with a phase stabilization occurring specifically between 8 and 12Hz and lasting 300ms (Fig. 4d). This absence of difference in CS− evoked freezing mechanisms is consistent with our behavioural observation of similar levels of CS+ evoked freezing (Fig. 3f).

**Figure 04:**
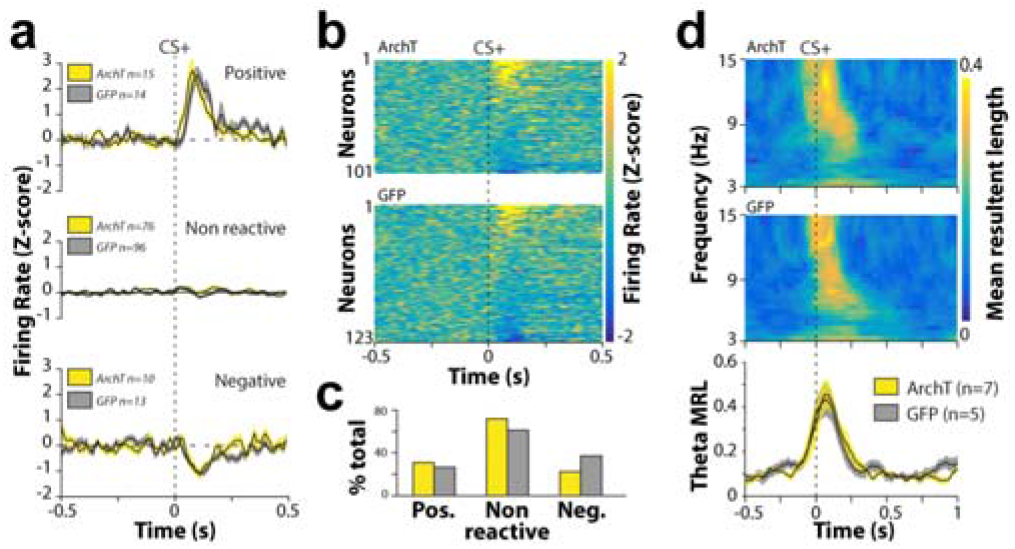
dmPFC coding mechanisms of conditioned stimulus is independent from previous dopaminergic modulation. a. Average PSTH for dmPFC neurons recorded in ArchT and GFP animals. Average firing rate (z-scored normalized according to pre-CS+ epoch) −0.5 to 0.5s before and after CS+ occurrence in excited (top), non-reactive (middle) and inhibited neurons (bottom). b. Heat-map displaying individual neuron firing behaviour in ArchT and GFP animals. c. Proportion of neurons excited, nonreactive and inhibited neurons after CS+ presentation (Khi-square test, X^2^(2) = 3.883, p = 0.7). d. Dopaminergic silencing during learning, does not affect dmPFC theta phase resetting to CS+ during memory test. Top. Mean resultant length (MRL) measures of phase stability across multiple CS+ presentations. The heat maps show an increase in MRL/stability between 8 and 15 Hz after CS+ presentation indicating that theta rhythm phase (8-12Hz) resets after CS+. Bottom. Average 8-12Hz MRL across animals shows that phase resetting is similar in ArchT and GFP groups. Shaded areas: mean +/− SEM

#### dmPFC neuronal synchronization and oscillations during freezing depend on prior VTA inputs

We then investigated whether dmPFC mechanisms of internally triggered/sustained fear expression had been impacted by prior VTA inputs suppression. These mechanisms consist in the oscillatory synchronization of dmPFC neurons on 4Hz (3-6Hz band) LFP waves^15^. We first measured the level of synchronization between dmPFC neurons within CS+, post-CS+ and within freezing in general by quantifying the amount of PN pairs that were significantly co-activated in each animals of ArchT and GFP groups (Fig. 5a,b). On average ArchT animals presented a smaller amount of co-activated pairs in post-CS and freezing epochs, but not within CS (Fig 2b). We then analyzed 4Hz rhythm and the relationship between 4Hz and dmPFC PN in both groups. Power spectral density computed between 3 and 16 Hz were highly similar in both groups with similar density distributions and average power in the 3 to 6Hz frequency band (Fig. 5c). Since recent studies have highlighted the olfactory system as the generator of this 4 Hz rhythm, it is but logic that these oscillations remained unaffected by VTA manipulations. Although this rhythm is conserved, the spike-phase relationship was significantly altered in ArchT animals. The analysis of 4 Hz phase-locked PN revealed a quantitative decrease for freezing epochs while no difference could be detected during CS+ (Fig. 5d). The deficit in freezing observed outside CS+ epochs (Fig. 3g) could hence stem from this deficit in neuronal synchronization and oscillations. Together with our behavioural observations, this show that DA inputs during conditioning contribute to the oscillatory synchronization that characterizes dmPFC PN during associative fear memory expression^14^.

**Figure 05:**
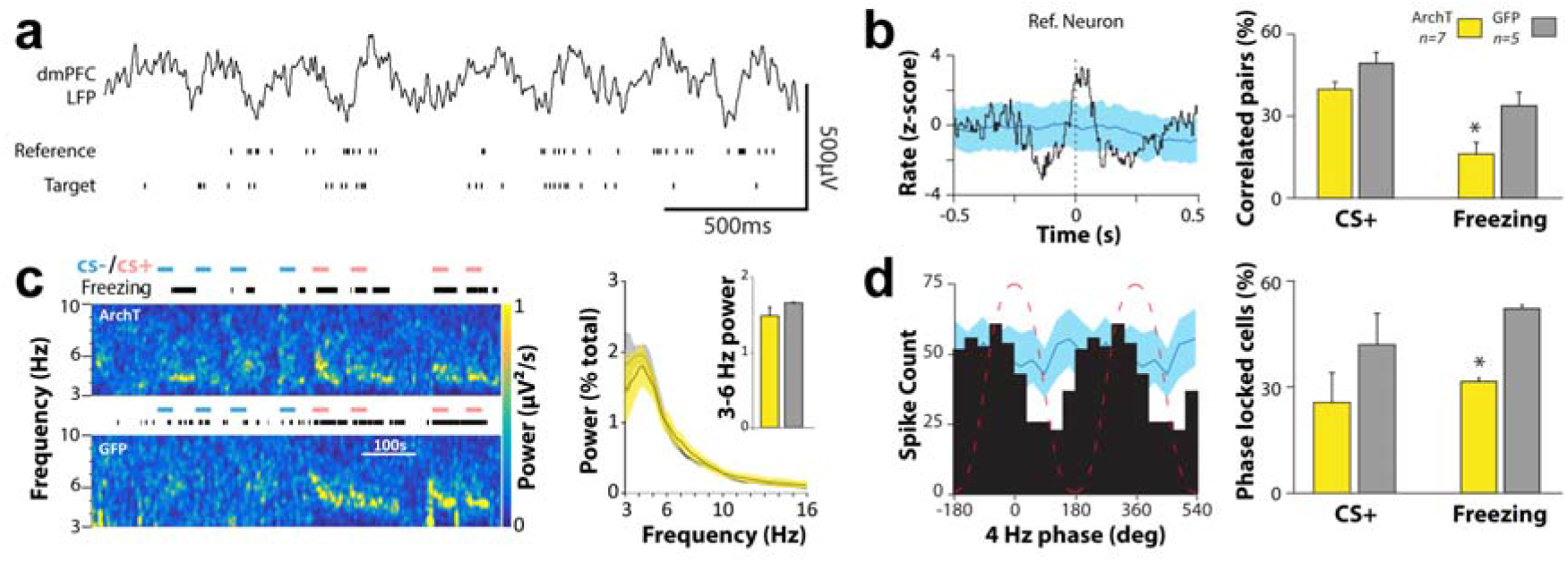
Inhibition of dopaminergic inputs during conditioning reduces dmPFC neuronal synchronization and oscillations outside CS+ presentations. a. Simultaneous recording of dmPFC LFP and single neurons showing the spike trains of a reference neuron and target neuron with which it will be compared for correlation. b. Left. Example crosscorrelogram (Z-scored) between the pair of neurons shown in a (blue area: 95% confidence interval computed from a x25 shuffled surrogate dataset). Only those pairs of neurons with CC crossing the upper limit of the surrogate confidence interval were tagged as significantly synchronized. Right. Proportion of correlated PN pairs recorded during CS+, post CS+ and Freezing epochs for ArchT and GFP groups. The proportion of ArchT dmPFC PNs correlated during post CS+ and freezing was significantly lower than in GFP animals. (ArchT n = 7, GFP n = 5, Mann-Whitney test, CS+: U = 6.00, p = 0.073; Freezing: U = 3.00, p = 0.018). c. Left. ArchT and GFP example spectrograms during test sessions depict power variations of dmPFC LFP oscillation as a function of time, frequency and freezing behaviour. Note that initial onset of 4 Hz oscillations is in the upper reaches of the 3 – 6 Hz range and, just as freezing, occurs inside and outside CS+ presentations. Right. Power spectral density histograms of dmPFC LFP during freezing. Shaded area: SEM. Inset: Comparison of 3-6 Hz average power during freezing between ArchT and GFP groups. No significant difference was found between the two groups (Mann-Whitney test, U = 10, p = 0.268). Error bars: SEM. d. Left. Example phase histogram for a neuron significantly phase-locked to 4Hz during freezing. LFP phase is represented by the dotted red line. Bin size 36°, 95% confidence interval derived from x25 shuffled surrogate dataset. Only spike histograms falling outside the confidence limit were tagged as significantly phase-locked and used for further analysis. Right. Proportions phase locked PNs in ArchT and GFP groups during CS+, post-CS+ and freezing (ArchT n = 7, GFP n = 5, Mann-Whitney test, CS+: U = 12.00, p = 0.432; Freezing: U = 5.00, *p = 0.048).

#### Dopaminergic inputs during fear conditioning control the precision of dmPFC ensemble coding of fear

Moreover, this transient, oscillatory synchronization in dmPFC PN population underpins specific network states that are necessary for the expression of freezing behavior, whether it occurs during or outside CS+ presentation^14^. Hence we investigated the impact of DA manipulation during conditioning on this crucial type of coding. To this end, we first computed the time resolved cross-correlation matrix between all simultaneously recorded PNs. This multidimensional set of matrices was then reduced by principal component (PC) analysis, where the first PC score captures dmPFC network state (NS) as a function of time (Fig. 6a). We hypothesized that the deficit in oscillatory synchronization in ArchT animals should impair the strength of population coding in ArchT animals and test this by measuring the distance between NS during freezing and non-freezing epochs (fig. 6b). While control animals showed a significant segregation between fear and non-fear NS distributions, the former were largely overlapping in ArchT animals and as a consequence, the Mahalanobis distance was significantly smaller in ArchT animals compared to GFP control. Finally we interrogated the decoding strength of dmPFC NS. In other words, we measured the accuracy by which one could retrieve animal behaviour just by looking at network activity (Fig. 6c). In both groups, NS predicted behavior with a higher-than-chance accuracy (chance level computed through a x100 shuffled surrogate data). Nonetheless, freezing prediction accuracy was weak in ArchT (57%) and significantly lower than that in control conditions. These results highlight the role of dopamine inputs during fear condition by enhancing the temporal precision and the accuracy of associative fear coding by neuronal population of the dmPFC.

**Figure 06:**
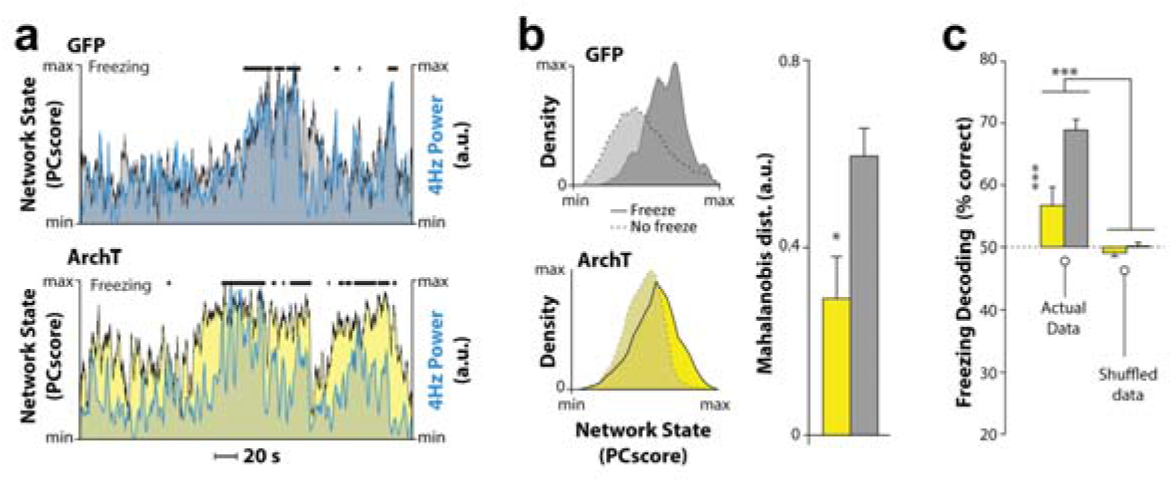
Dopaminergic inputs support the precision of dmPFC ensemble coding of fear. a. Joint time course of dmPFC neuronal population states, 4Hz oscillations and freezing behavior. Examples of GFP (top) and ArchT (bottom) network state as a function of behavior. Network state is measured as the principal component score of single unit crosscorrelation matrix and captures the different synchronization states of dmPFC neuronal network. In this examples, high PC scores and corresponding network states co-occur with both freezing and increases in LFP 4Hz (blue line). b. dmPFC in ArchT animals presents a deficit in fear state discrimination. Right. Examples of GFP (top) and ArchT (bottom) network states distributions for freezing and non-freezing epochs. Note the strong overlap between distributions in ArchT animals. Left. Mahalanobis distance between freezing and non-freezing network states is significantly reduced in ArchT animals, indicating a lack of network discrimination compared to GFP controls (ArchT n = 5, GFP n = 5, Mann-Whitney test, U = 2.00, p = 0.03). c. Freezing prediction from dmPFC network activity is weakened by priorz dopaminergic silencing. Freezing decoding from neuronal ensemble activity present a modest accuracy (57% of correct prediction) and is strongly decreased compared to GFP controls (69%). Note it is still possible to decode animal’s behavior in both group as shown by a significant difference between actual and shuffled dataset (ArchT n = 5, GFP n = 5, Two-way ANOVA with one factor repetition, Factor 1: ArchT vs GFP, F(1,8) = 15.933, p = 0.004; Factor 2: data vs. surrogate, F(1,8) = 50.942, p < 0.001; F1×F2, F(1,1) = 9.363, p = 0.016; Holm-Sidak post-hoc adjusted multiple comparison: within data, ***p < 0.001).

## Discussion

### VTA coding of conditioned stimulus

Our data unravel that 2 different types of coding develop for through consecutive pairings of CS+ to the shock. The majority of both rate and signal-to-noise coders maintained their coding of CS+ during memory testing and also displayed a dip in firing right after CS+ termination when US is supposed to occur. These features remind of the US to CS transfer of dopaminergic activity theorized by Rescorla-Wagner^13^ and its experimental counterpart^7^, hence extending the former to associative learning driven by stimuli of negative valence. During fear acquisition, these coding neurons also preferentially synchronize with dmPFC 4 Hz oscillations. Both Rate and SNR coders likely provide a timely and transient increase in dopamine within the dmPFC that coincides with the excitation of local neurons by the stimulus. This process is prone to synaptic long term potentiation in and between cortical neurons^20^ that supports the formation of functional population ^21^, especially in recurrent networks such as the neocortex ^22^.

### Dopamine role in dmPFC coding of fear behaviour

Shutting down VTA dopaminergic inputs during CS− US association translated into a deficit in internally triggered fear behaviour indicating that VTA-dmPFC synchronization is important for this aspect of fear memory neurophysiology and particularly local cortical mechanisms. In dmPFC, population coding is necessary for the expression of learned fear, whether the memory is reactivated by a CS^13^ or internally^14^. In our study, CS induced freezing and mechanisms remained unaffected by prior dopaminergic silencing. This might derive from at least two reasons. One, manipulation of the terminals severs information flux towards the cortex but do not prevent DA neurons to gain associative coding properties that might arise when the stimulus is presented. This is supported by recent results showing that acute modulation of VTA inputs during memory retrieval affects dmPFC function together with fear expression^12^. Another non-exclusive hypothesis is that CS induced coding might depend on other neuronal elements such as the VTA to BLA pathway^23^. The main effect we observed is on internally triggered fear (outside CS presentation). VTA inhibition during conditioning results in a decrease of the frequency of those episodes and alterations in dmPFC population dynamics. The transitions between non-freezing and freezing network states are less marked and this correlates, and likely translate, with a lower probability of freezing initiation. Besides these transition epochs, on a more global scale there is little difference between freezing and non-freezing network states after DA manipulation and a poor prediction accuracy regarding behaviour. This can be interpreted as a lower signal to noise ratio for structures driven by dmPFC such as the amygdala or the periaqueductal grey. Dopamine enhances neuronal gain to synaptic inputs and is therefore a strengthening element for functional neuronal assemblies. Under dopamine depletion, cortical attractor networks have been theorized to be less stable^24^. Our data support this view in that the sound neuronal discrimination between behavioural states in control conditions (deep stable attractor) turns into a poorly relevant predictor when the network as settled its functional pattern in the absence of dopamine (shallow unstable attractor).

Although not absolutely necessary to fear memory expression, dopamine as a neuromodulator is instead a stabilizing element for neuronal assemblies controlling fear expression. From a therapeutical standpoint, it has been suggested that dopaminergic dysfunction could participate to PSTD development^25^. Our results support this idea and the use of dopamine antagonists could be envisaged as a potential complement of current therapeutic approaches of fear and anxiety related disorders.

## Acknowledgement

The authors thank members of the Herry lab for discussion and constructive comments on the manuscript. This work was funded by ANR Programme Blanc.

---

**Supplementary figure 1:**
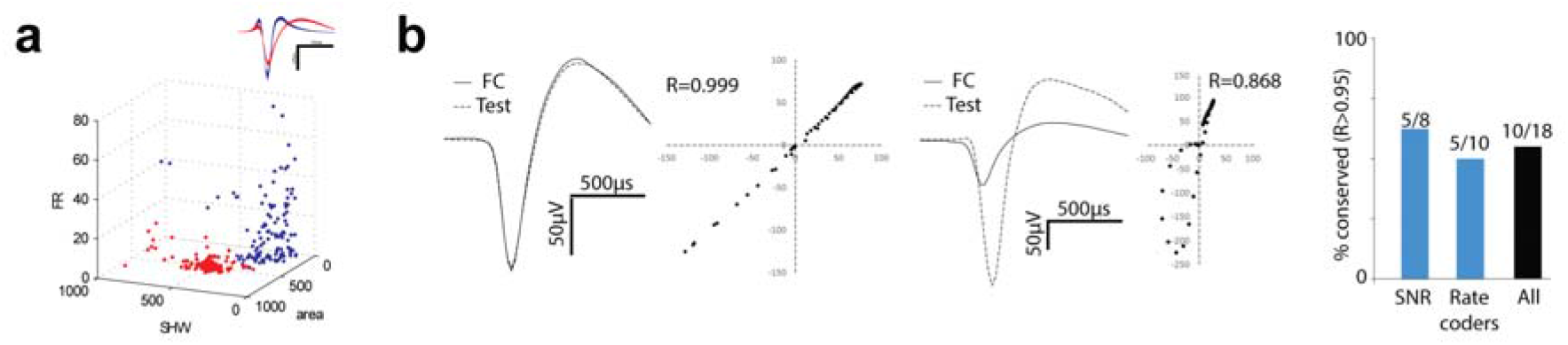
Single neuron characterization. a. Cell type discrimination. Putative principal neurons and interneuron clustering (red, interneurons - blue) have been identified based on 3 biophysical criteria; firing rate, spike half width (SHW) and area under the curve (AUC) of the waveform. b-c. Waveform stability analysis. b. left. Example of a stable unit’s recorded waveform across Fear conditioning (FC) and Test recording sessions. Right. Unstable unit example. c. Proportions of stable units (R>0.95) conserved along FC and test sessions among signal to noise ratio coders (SNR), rate coders or both (All).

**Supplementary figure 2:**
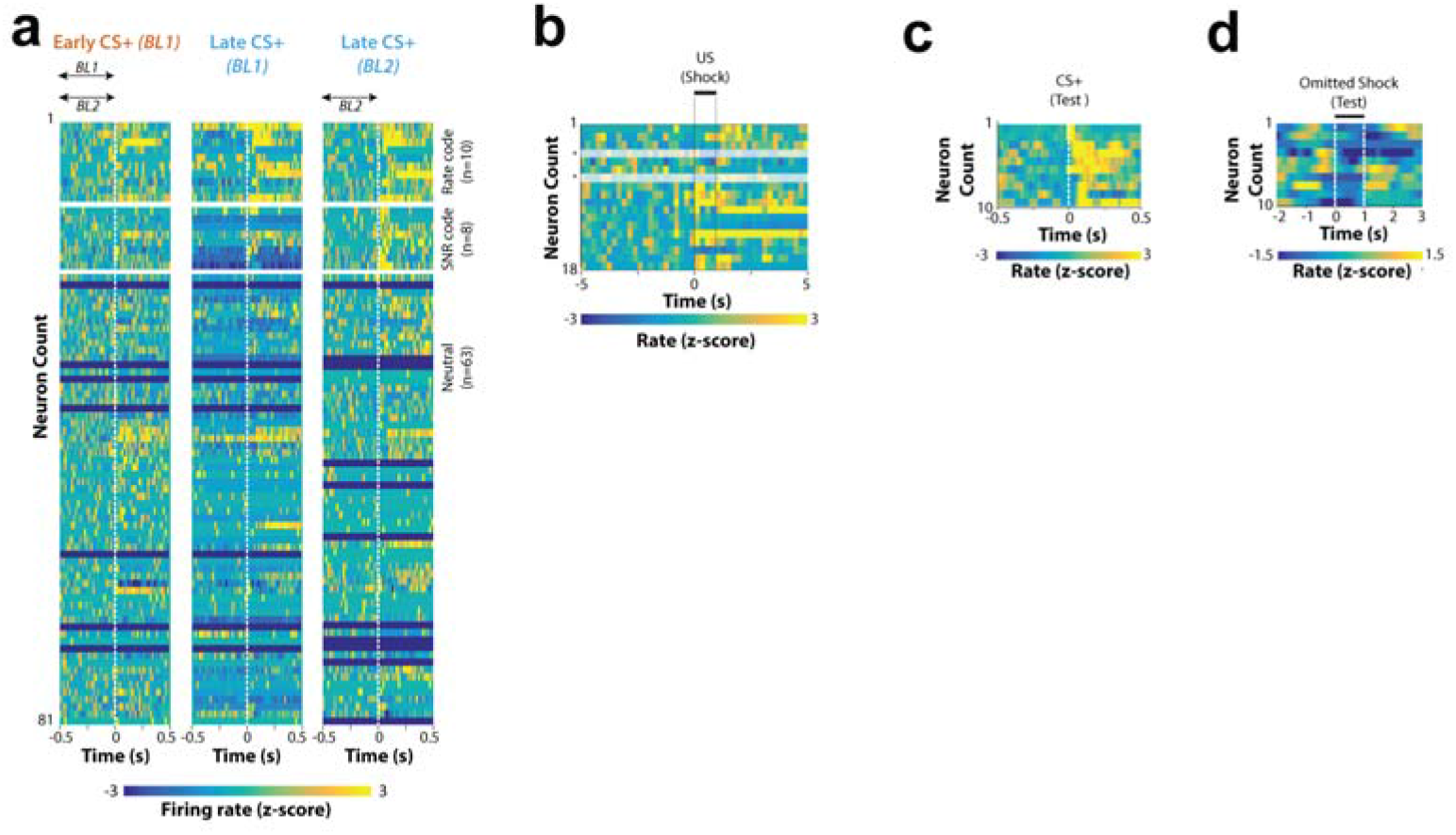
a. PSTH centered on CS+ for all 81 neurons recorded during the fear conditioning. From left to right: Early CS+ response normalized onto early pre-CS+ (Baseline 1 BL1), Late CS+ response normalized onto early pre-CS+ (BL1), Late CS+ response normalized onto late pre-CS+ (BL2). b. PSTH centered on US (Shock) for all 18 coding neurons. Two of those where omitted because the electrode they were recording from presented signal loss (clipping) during and after US delivery. Two of those where omitted because the electrode they were recording from presented signal loss (clipping) during and after US delivery. c. PSTH centered on CS+ during test for the 10 coding neurons that were recorded over FC and Test days. d. PSTH centered on shock omission moment during test for the same 10 coding neurons.

**Supplementary figure 3.**
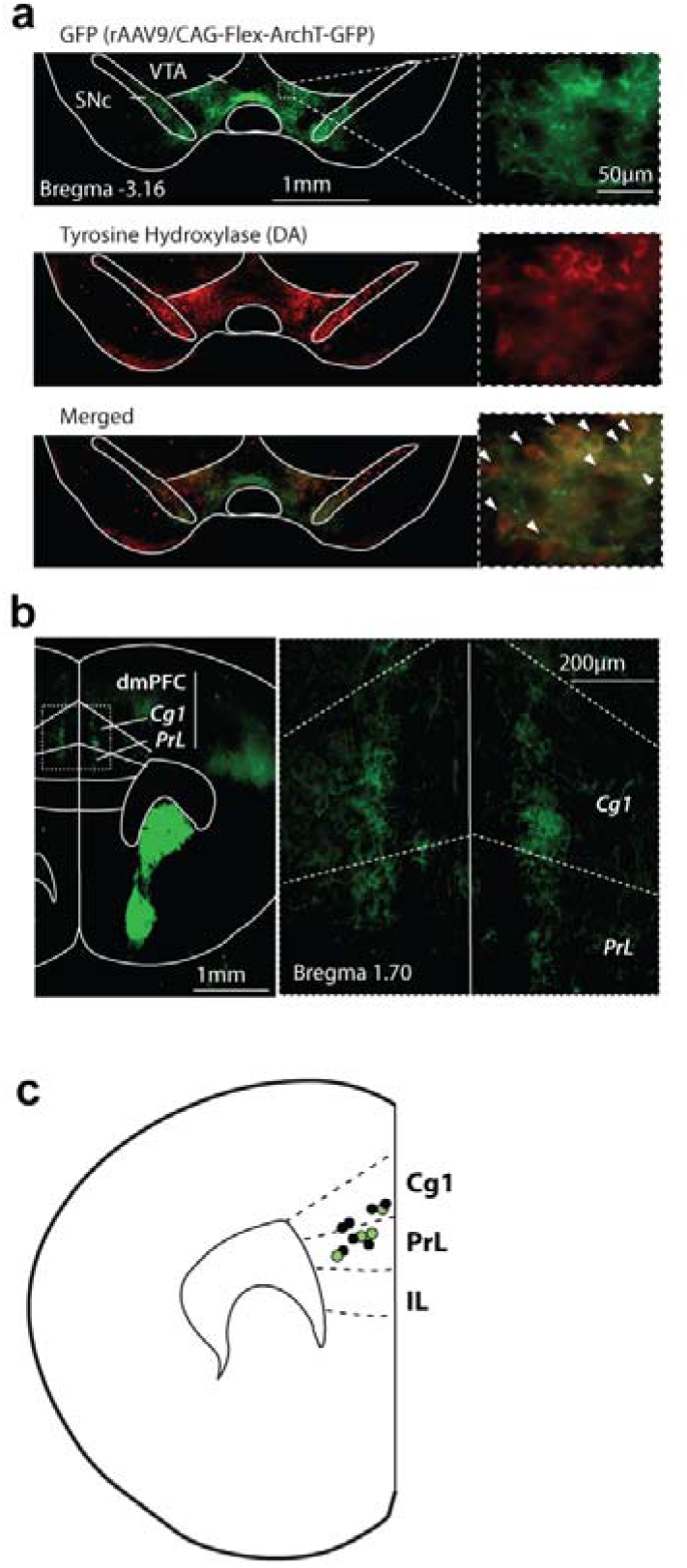
a.b GFP / tyrosine hydroxylase immunohistochemistry in DAT-Cre mouse brain. GFP and tyrosine hydroxylase (TH) immunostained brain sections imaged with epifluorescent microscope displaying transfection of VTA DA neurons and projection to dmPFC. a. Top : GFP immunostained (Alexa Fluor 488) section labelling expression of Cre-dependent ArchT expressing AAV virus in the VTA. middle: Red fluorescent dye (Alexa Fluor 594) labelling for TH. Bottom : overlay of ArchT and TH expression. Arrows identify overlap of ArchT and TH positive cell bodies. b. GFP immunostained brain section showing structures receiving afferent DA projections from the VTA. Enlargement (X 10) shows GFP expression in DA neuron terminals in the dmPFC (Cg1 & PL). c. Optic fiber and optrode/electrode implant sites recovered from PFC brain sections (yellow: ArchT animals, gray: GFP animals). Cg1 – cingulate cortex; IL – infralimbic cortex; PrL – prelimbic cortex; SNc – substantia nigra pars compacta; VTA – ventral tegmental area.

**Supplementary Figure 4:**
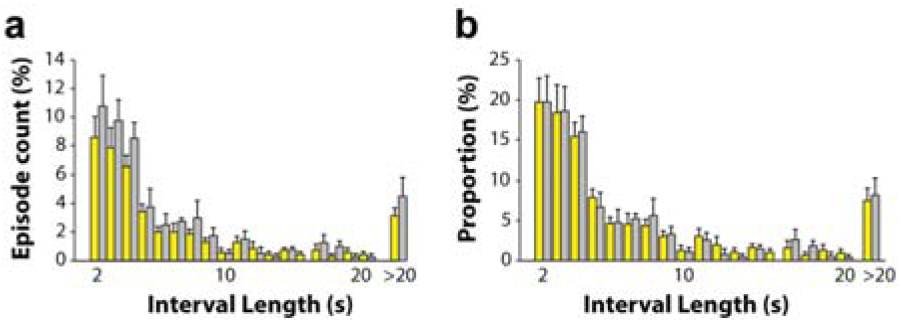
Distribution of fear episodes as a function of their length. a. Absolute episode count. b. Proportion relative to total episode count (Yellow: ArchT, Grey: GFP)

## Notes

### Competing Interest Statement

The authors have declared no competing interest.

